# Inversions and integrations using CRISPR/Cas9 and a module-based plasmid construction kit for fission yeast

**DOI:** 10.1101/2023.10.02.560446

**Authors:** Cristina Berenguer Millanes, Bart P.S. Nieuwenhuis

## Abstract

Structural variants (SV) are genetic alterations that involve large-scale changes in the structure of a genome. These variations can encompass deletions, duplications, inversions, translocations, or complex rearrangements. While smaller structural variants are relatively well studied, much is unknown about the prevalence and effect of larger SV. Genome sequencing methods that are able to find large reorganization in the genome, e.g. using mate pair or long-reads, have shown inversions and translocations are more common than was previously expected. Reduced recombination between regions with different structural organizations leads to the rise of variant specific alleles. Studying the effect of the SV in isolation is obscured by their independent evolutionary histories. Tools are needed to introduce SVs without introducing correlated alleles. Here we describe a method to introduce specific inversions and rearrangements in the fission yeast *Schizosacchromyces pombe* using the modified CRISPR/Cas9 system SpEDIT to introduce multiple breakpoints with a single plasmid. Sequences for homologous recombination that guide repair resulting in the desired SVs are generated using an extended method for Golden Gate DNA shuffling for *S. pombe*. Our extension of the system from Kakui et al. is more efficient for integration, introduces more flexibility, and extends the system beyond single construct integrations. Additionally, we extend the set of promoters, tags, markers and terminators, specifically using DNA sequences from other fission yeast species, which avoid introduction of homologous sequences, thereby reducing the chance of non-allelic homologous recombination during sexual reproduction.

## Introduction

Recent research has shown that the structure of the genome has many implications on complex phenotypic characteristics (Weischenfeldt et al. 2013; Gorkovskiy and Verstrepen 2021). Inversions, translocations, and chromosome fusions can have effects at the whole genome level, leading to genomic instability and chromosomal imbalance after meiosis. Also at the genic level, SVs have important consequences. Rearrangements can result in genes being positioned in new genomic contexts, which can affect their expression patterns (Lavington and Kern 2017; Huang et al. 2018). Structural variants can affect regulatory elements, which can lead to dysregulated gene expression patterns, potentially affecting developmental processes and cellular functions. Furthermore, SVs can reduce recombination, greatly affecting adaptation and can even lead to speciation (Kirkpatrick 2010; Thompson and Jiggins 2014). To study these types of effects, efficient manipulation of the genome structure is required. Here we present methods for the fission yeast *Schizosaccharomyces pombe*, to efficiently introduce translocations and inversions, and introduce genetic constructs at many locations in the genome. Our method combines CRISPR/Cas9 with homologous recombination guides generated by a flexible Golden Gate method that can be used for efficient targeting and to generate structural variants (SV) in the genome.

A variety of tools has been used over the last decades to manipulate the genomic structure in fission yeast. Reorganizations of large regions of the genome can be performed using Cre-loxP (Ishii et al. 2008; Hu et al. 2015; Gu et al. 2022) and site specific HO endonuclease systems (Sunder et al. 2012). These systems have yielded great insights, nevertheless they are not very flexible, and they require introduction of genetic elements into the genome or introduce scars at the sites of manipulation. Since the introduction of CRISPR/Cas9 molecular manipulation of the nuclear genome has become very efficient. The enzymatic induction of directed double strand breaks has greatly simplified genomic modifications and has increased efficiency of genetic modifications in the genome (Hsu et al. 2014). Together with introduction of sequences that induce homologous recombination in the regions flanking the double strand break, deletions and integrations can be made with high efficiency (Rodríguez-López et al. 2017).

Introduction of genic elements at specific locations in the fission yeast genome has been well established since the 1970s, however these were often depend on laborious cloning strategies. A few decades later, PCR based methods for integration of specific sequences at any position were developed (Bähler et al. 1998; Snaith et al. 2005). These methods, though highly flexible in changing the target, have a few drawbacks. First, introduction of genetic elements this way can introduce unwanted variants due to PCR errors and second, due to relatively short regions of homology, they are not very efficient and can be promiscuous (Murray et al. 2016). Even though these plasmids are flexible in the target of integrations, they are limited in the elements that are to be integrated. Other plasmid tools that are more flexible exist in which a gene of interest, a tag or fluorescent marker can be easily integrated (e.g. Kakui et al. 2015; Vještica et al. 2020). However, these often are targeted to specific genes such as the *ura4*, *ade6*, or *his5* and lack the flexibility to integrate anywhere into the genome. We present a flexible Golden Gate based system that can be used for efficient construction of HR fragments and be targeted to any region of the genome.

Golden Gate Assembly is a powerful molecular cloning method that allows the efficient and precise assembly of multiple DNA fragments into a larger DNA construct in a single reaction. The method employs type IIS restriction enzymes, which cleave DNA outside their recognition sequence, generating unique overhangs. These overhangs act as cohesive ends that can be specifically ligated with complementary overhangs from other DNA fragments. We build on the system initiated by Kakui et al. (2015) with extension by Kiriya et al. (2017) to generate stable integration fragments that can be introduced into heterologous regions of the gnome. The plasmids avoid integration of repetitive sequences using single integration sequences (Vještica et al. 2020) and avoid integration of plasmid elements by eliminating the backbone from the integrated fragment (Bähler et al. 1998). Inspired by the Golden Gate system from Binder et al. (2014) we added a lower *BbsI* level to the system which allows for easy generation of homologous integration locations and a higher level, that allows for easy assembly of multiple elements. Additionally, we have extended the set of available elements with Promoters, Terminators and Fluorescent tags. To avoid repeated homology with other sites in the genome, the promoters are obtained from fission yeast species other than *S. pombe* and terminators from *Saccharomyces* species. We provide a variety of targets for integration locations each with a CRISPR/Cas9 plasmid with the single guide RNA (sgRNA) element that directs to the integration site. However, the modularity of the Golden Gate system is easily expandable and highly flexible to generate novel targets.

To generate structural variants, two sgRNA are required at the same moment, to generate the pair of DSB that are the break points for rearrangements. We modified the Golden Gate method from Torres-Garcia et al. (2020), in which a single guide is introduced in a plasmid that contains the Cas9 gene and a selective marker. We designed an extra fragment that can be used with the plasmids from Torres-Garcia et al. to integrate two guides into a single plasmid in a single reaction. We present these methods and give three examples of inversions that were generated using this method.

## Results

We present an extended method to use CRISPR/Cas9 combined with Golden Gate (**Fig. 1**), to efficiently introduce genomic elements into the fission yeast genome, and to introduce genomic rearrangements and inversions. Note we only describe the system for C-terminal tagging, but that N-terminal tagging is possible too (for details see Kakui et al. 2015). We start with presenting the elements of the assembly method cloned in *Escherichia coli*, followed by presenting results for the integrations and SVs generated using these methods in fission yeast.

**Figure 1.**
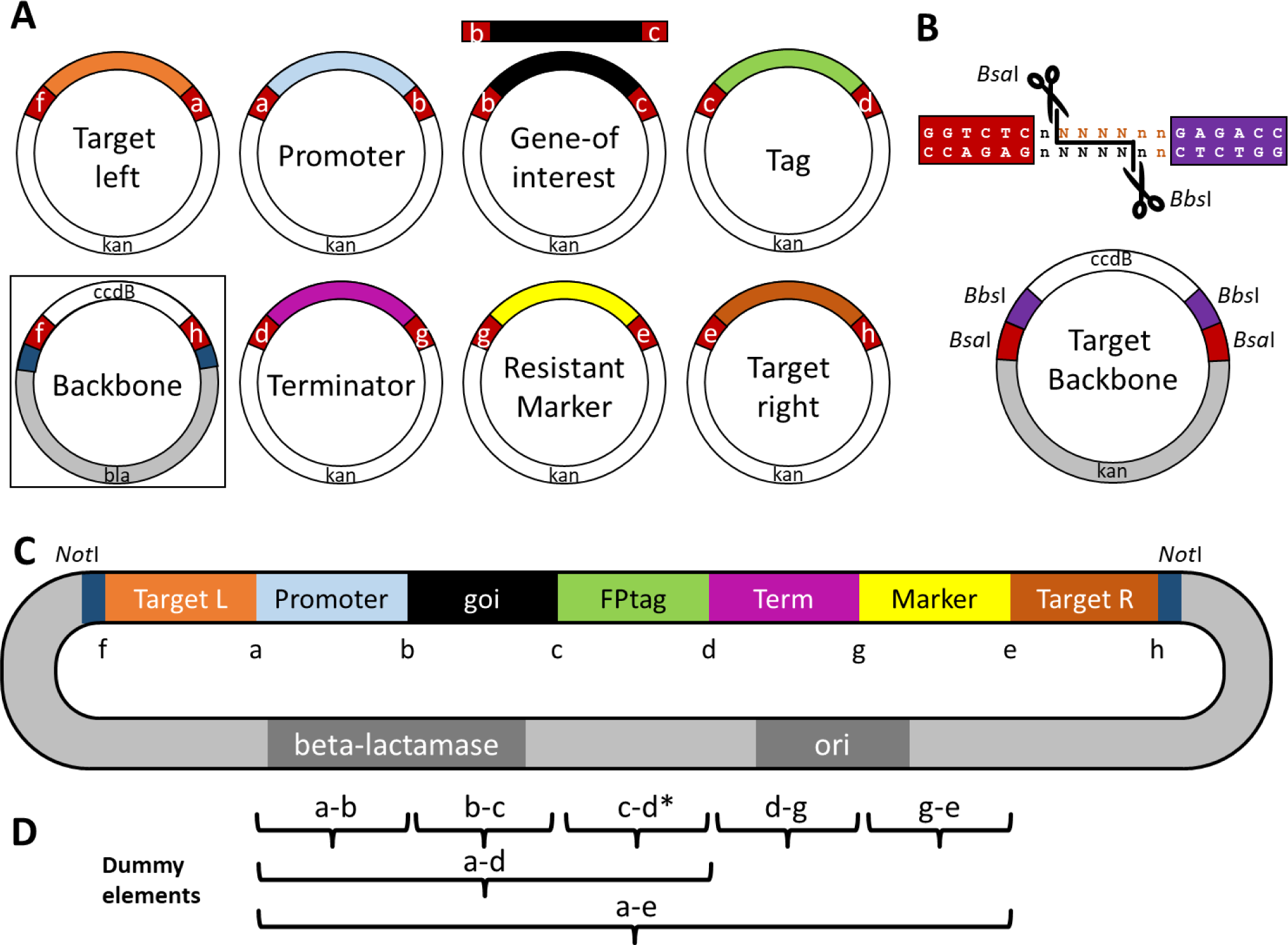
Overview of the golden gate elements and contruction of level 1 and level 2 elements. **A** The donor elements with overhangs are indicated by letters and the white parts contain the BsaI restriction sites, which will not be present in the final product. Each donor contains a kanamycin resistance gene in the backbone, except for the (f-h) backbone which is ampicillin resistance. This latter plasmid additionally contains a ccdB gene that kills all cells containing undigested backbone plasmids. The (b-c) and (c-d) elements can be reversed for N-terminal tagging as explained in Kakui et al. (2015) **B** Plasmids for Level 1 target elements contain *BbsI* recognition sites that are cut when generating L1 donor plasmids, which can be used to generate L2 plasmids utilizing the *BsaI* sites in the backbone. Additionally, they contain a ccdB that is excised during assembly. **C** Example of a fully constructed Level 2 plasmid. The to be inserted fragment can be excised from the backbone using *NotI* digestion and directly used. Alternatively, the fragment can be amplified by PCR using primers M13F and M13R. **D** A set of dummy elements has been produced to span gaps that are not required in an assembly.

### Extended Golden gate assemblies

The Golden Gate Assembly method assembles multiple fragments in a single reaction, either for integration or for epitopic expression (**Fig. 2**). DNA fragments that are either generated by PCR or are present in a donor plasmid (Level 1) are digested with type IIS restriction enzymes and ligated (Marillonnet and Grützner 2020). The digested DNA fragments are designed to have complementary overhangs that can assemble in one specific order into a larger DNA construct resulting in Level 2 plasmids. After transformation into *E. coli*, only cells containing the desired construct are able to grow due to different antibiotic resistance genes in the donor and recipient vector. Level 2 plasmids can be combined in a next assembly step (**Fig. 2**) resulting in epitopic expression plasmids (Level 3), that can be modified for genomic integration in a final assembly step (Level 4).

**Figure 2.**
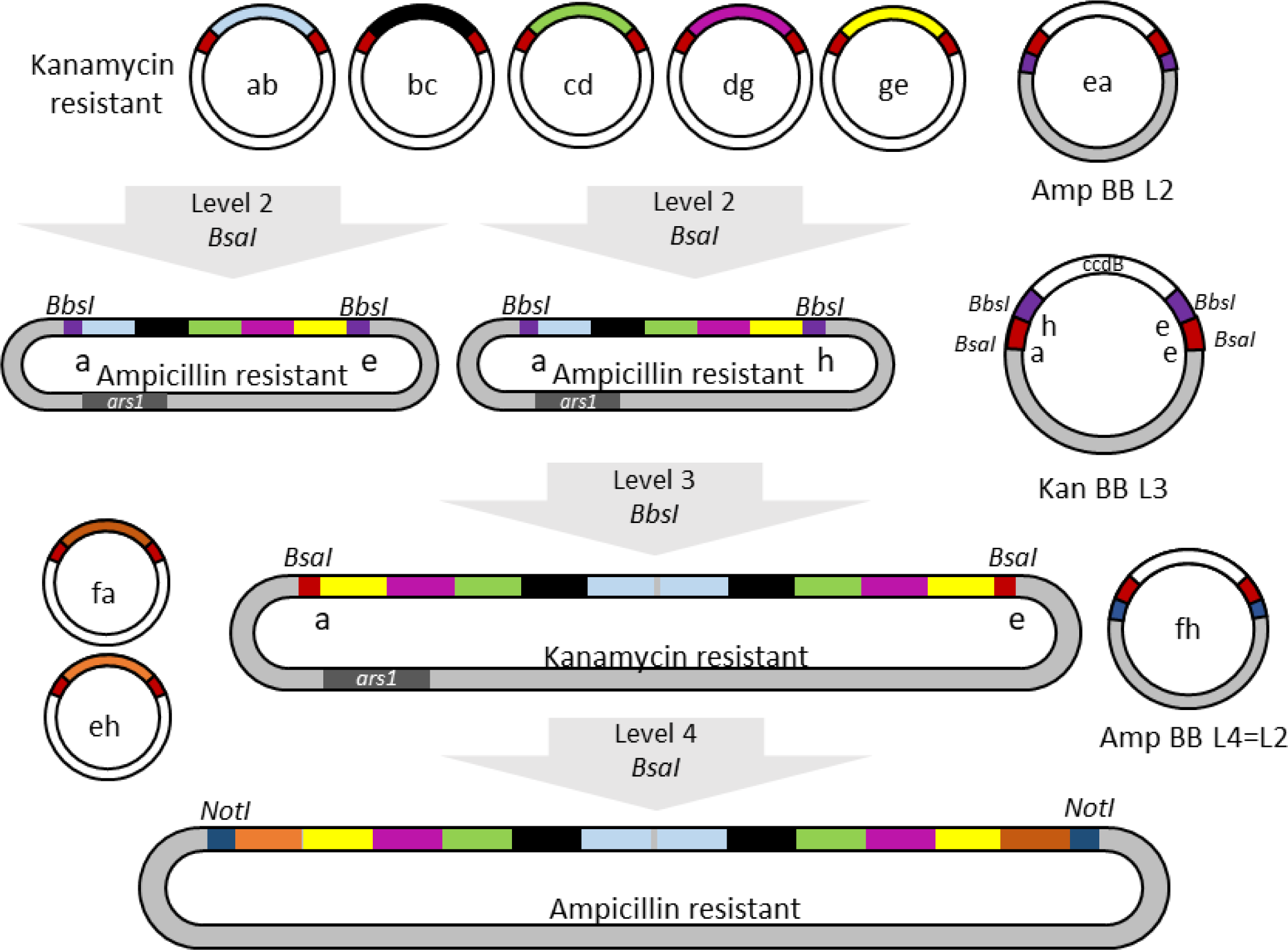
Overview of the extension of the golden gate system that allows for epitopic expression (L2) and combination of two constructs for epitopic expression (L3) or integrations in a single location (L4). Kanamycin resistant elements are assembled using *BsaI* as in Fig. 1 now using an ampicillin backbone with (e-a) overhangs that contains an *ars1* sequence. These L2 plasmids can be directly used for epitopic expression or two with complementary overhangs can be assembled together into an L3 plasmid using *BpiI* and selecting for kanamycin resistance. In a final step, this expression vector can be changed into a double integration element by adding the target regions from L1 plasmids using *BsaI* and selecting for ampicillin resistance like for an L2 assembly.

#### Donor elements (Level 1)

Each element’s position in the final construct depends on the overhangs of the *Bsa*I restriction sites. After choosing the correct overhangs, primers are designed with tails that contain the *Bsa*I sequence and the four bases for the Golden Gate system. We extended the existing system of overhangs with two sequences for newly added homologous recombination elements (Kakui et al. 2015; Kiriya et al. 2017). These elements allow for integration of the construct at any position in the genome. A full overview of the cut-sites and primer overhands can be found in **Fig. 1a** and **Supplementary Table 1.**

Instead of using expensive TOPO backbones for cloning (Kakui et al. 2015), we designed a cheap and efficient system to add elements to the Golden Gate set. We used a backbone in which we introduced a *Sma*I cut-site in the multi cloning site. This cut-site allows for introduction of blunt-end fragments in a single restriction-digestion step, similar to the blunt-end cut-ligation in Binder et al. (2014). Fragments to be integrated are generated using Phusion PCR or for short fragments directly from annealed primers. Successful integration disrupts expression of the toxin ccdB and, when using cells sensitive to ccdB (e.g. DH5α, TOP10), only cells with a successful integration plasmid will survive. Tests showed that most colonies that grew after integration contained a fragment of the correct length as confirmed by colony PCR. Initially we tested 8 colonies by PCR, but as almost all colonies were positive, we now standard only test 4. Due to the presence of *E. coli* promoters near the integration site, some genes might be hard to clone. For example, we were unable to clone into the backbone the *mam2* gene, which encodes a seven-transmembrane G protein-coupled receptor, probably due to toxicity of the protein in *E. coli*. Selected colonies were cultured and plasmids isolated with a miniprep. Due to blunt-end ligation the directionality of the insert cannot be predicted, but for L1 donor elements this is not relevant. We recommend sequencing of the margins using M13F and M13R primers, to assure presence of the *BsaI* restriction sites and the correct Golden Gate overhangs. When needed, a *BsaI* or *BbsI* restriction site can be removed using site directed mutagenesis of the assembled plasmids.

To manipulate the genome at a variety of sites by integration or to induce inversions, flanking regions with homologous sequences are required. To avoid errors in the overhangs and restriction sites for these elements, we created two plasmids for (f-a) or (e-h) overhangs, respectively. The homologous regions are amplified by Phusion PCR with primers with overhangs containing *BbsI* cut sites that are integrated into these plasmids (**Fig. 1b**). During the integration, the *ccdB* gene and the *BbsI* cut sites located in the receiving backbone are eliminated, while *BsaI* cut-sites in the backbone become available for assembly into level 2 plasmids (see below). This ensures a high efficiency of plasmid construction and avoids damage of the *BsaI* cut sites in these plasmids. Additionally, it facilitates assembly of multiple elements using *BbsI* to remove unwanted cut sites (Binder et al. 2014). For example, targets to integrate fragments directly after the *mat1P* or *mat1M* were generated in this way. Two or three fragments respectively, were amplified using primers with a synonymous mutation at *BsaI* sites, each with a *BbsI* overhang that were assembled in a single reaction into the (e-h) backbone (**Fig. S1**). The result is a donor fragments without *BsaI* or *BbsI* sites that can be used for Golden Gate assembly.

#### Golden gate assembly Levels 2, 3 and 4

Assembly of the elements into a single construct is performed in a single reaction in which all fragments need exactly one other fragment with a complementary overhang on either side. The full construct can be assembled from PCR fragments or the Level 1 plasmids into a Level 2 plasmid. The resulting plasmid combines the elements in the following order (overhangs in parenthesis; **Fig. 1c**): left homologous region (f-a), promoter (a-b), gene-of-interest goi (b-c), tag or fluorescent marker (c-d), terminator (d-g), selectable marker (g-e) and right homologous region (e-h). The elements are introduced into backbone pBN411 using the overhangs (h-f). Transformed *E. coli* are selected on ampicillin plates, which selects against the Level 1 plasmids, which are kanamycin resistant. The receiving backbone contains a *ccdB* gene, which is removed in successful assemblies, enabling selection against undigested receiver plasmid.

Not always all elements are required in an assembly. We generated a set of dummy elements, very short sequences to span gaps, which greatly improves flexibility and reusability of donor elements (**Fig. 1d**). Most dummies are short filler sequences without function, however dummy (b-c), which replaces the goi, contains an ATG and ends in-frame, and dummy element (c-d), which replaces the tag, has a stop codon in every frame.

Next to the integration vector, we produced a episomal expression vector with the yeast autonomous replication sequence *ars1* (Heyer et al. 1986) and (a-e) overhangs. Constructions of these plasmids happen similar to integration vectors, with plasmids selected on ampicillin plates. The constructs can be used for episomal gene expression similar to pBMod-exv from (Kiriya et al. 2017). We generated two version of this backbone (pBN212 and pBN231) that contain different *BbsI* restriction sites in the assemblies. These sites can be used to combine the elements generated in each of the plasmid and assemble them together in a Level 3 construct (using backbone pBN432 and selecting on kanamycin; **Fig. 2**). This L3 plasmid can be used as an expression vector in which multiple constructs can be introduced into the same cell in a single transformation event using single selection marker. Additionally, the L3 plasmid contains *Bsa*I restriction sites with (ae) overhangs and can be used as donor plasmid for Level 4 assemblies. In this final step, target regions are added to allow for integration of the assembled fragment into the yeast genome (using Level 2 backbone pBN411 and selection on ampicillin). The modularity of the system makes it possible to combine multiple markers easily, each with their own promoter, tag and potentially selective marker, at a single integration position in a single transformation event.

Assembly of each of the levels is very efficient, generally generating thousands of successful assemblies. Where we initially tested assemblies by colony PCR using primers over multiple fragments, since switching to BsaI-HF-v2 from NEB, the success rate has improved considerably and now we immediately perform minipreps. Verification by RFLP shows that for most contructs the assemblies are correct.

#### Chromosomal integration of elements

The HR fragments that were assembled as described above were introduced into the fission yeast genome using *CRISPR/Cas9*. Linear fragments were isolated from the backbone using either Phusion PCR or by restriction digestion using the *NotI* cut sites present in the backbone (**Fig. 1c**). Integration was performed either with plasmids from CRISPR4P (Rodríguez-López et al. 2017) or SpEDIT (Torres-Garcia et al. 2020) at a variety of sites, in all of the three chromosomes. There was no consistent difference in efficiency between CRISPR4P or SpEDIT plasmids, however we did see a large difference between the locations in the genome. Replacement inside CDS of genes, was most efficient, even more so when using genes resulting in prototrophic strains (e.g. *leu1, ade6, his5* etc.). We tested 14 intergenic regions that were completely without annotations in PomBase (Harris et al. 2022). Most of these had good transformation efficiencies, however, we were not successful in transforming two regions. Position II:2.29 never generated inserts in the desired location, and position II:2.20 generated transformants very rarely, and always resulting in diploid cells. Even though relatively long homologous fragments were used, tetrad dissections to assess single insertions showed, that two out of about forty transformation events double insertion had taken place. Verification for single insertions is thus advised.

### Non-*S. pombe* elements

Homologous genomic sequences at different loci in the genome can lead to non-allelelic homologous recombination (NAHR), which generates structural variation in the genome (Lambert et al. 2005).Integration of elements at different sites in the genome can be problematic when these contain identical or highly similar regions. This can be avoided by using sequences from other yeast species as shown by (Li et al. 2019). We followed their example and designed a set of promoters with (a-b) overhangs for which the *S. pombe* versions had been tested (Russell 1989; Iacovoni et al. 1999; Matsuyama et al. 2008; Watt et al. 2008; Verma et al. 2014; Wang et al. 2014). We derived these sequences from either *S. octosporus*, *S. japonicus* or *S. cryophylus*. (P*tif51*_so, P*cam1*_so, P*ef1a-c*_so, P*lsd90*_so, P*eno101*_so, P*inv1*_so, P*urg1*_sj, P*eis1*_sj, P*adh1*_sj, P*gpd3*_sj, P*mei2*_sc). Regions approximately 850bp upstream of the coding region were amplified.

We tested their efficiency by heterologous expression of CFP in *S. pombe*. Signal strength of CFP was measured by flow cytometry and compared to the P*adh1* from *S. pombe*. The strengths of the measurements after gating for singletons showed good expression for most. Most promoters were able to induce strong expression of the CFP, however, there was a clear difference in promoter strength. For promoters *Pcam1_*so and P*inv1_*so the results were inconclusive, as CFP did not appear induced. Promoters P*ef1a-c*_so and P*eis1_*sj showed medium strength, and the rest of promoters showed high to very high strength (**Fig. 3**).

**Figure 3.**
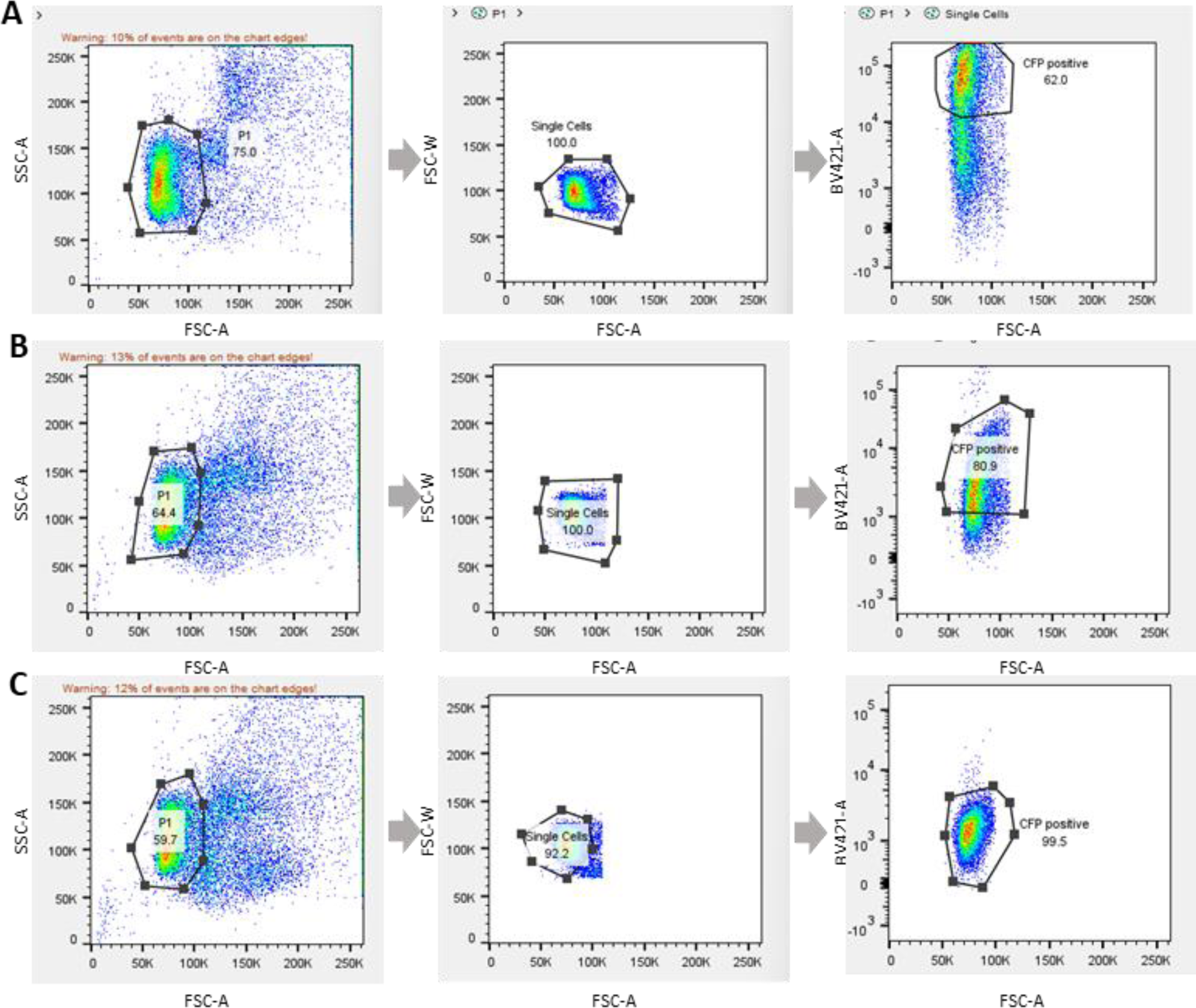
Examples of flow cytometry results for promoter strength analysed with FlowJo™ v10.9.0 Software (BD Life Sciences). In the three panels for both examples, from left to right, showing all events on the left with P1 preliminary gate, showing P1 selection with Single cells gate, and on the right showing the CFP positive hits. **A)** Example of expression for construct P*eno101_*so-CFP, a very strong constitutive promoter. **B)** Example for construct P*adh1_*sj-CFP, a strong constitutive promoter. **C)** Example for construct P*ef1a-c_*so-CFP, a medium strength constitutive promoter.

Additionally, we generated three auxotrophic markers derived from *S. octosporus* (*ura4, arg3* and *his3*) under their native promoter, which can be used to select for transformants in prototrophic *S. pombe* strains. The transformants were tested in epitopic expression vectors and were able to complement the prototrophic *S. pombe* strains (**Fig. S4**). Finally, we added four promoter regions to the collection of elements that are derived from different *Saccharomyces* species. These promoters have been used before in fission yeast (Li et al. 2019) and were added as (d-g) elements.

### Generating inversions and chromosomal rearrangements

Double strand breaks can be manipulated to promote repair in a desired way, by providing template DNA with segments for homologous repair, close to the break points. CRISPR/Cas9 in fission yeast has been efficiency used to delete segments (Rodríguez-López et al. 2017) introduce elements (Torres-Garcia et al. 2020) or introduce SNPs (Zhang et al. 2018). In all cases, a single guide RNA is used to induce a break in the genome that is repaired by an HR fragment. When multiple guides are introduced into the same cell, genomic shuffling is possible during which sequences flanking different breakpoints can be combined together (Fleiss et al. 2019).

We used CRISPR/Cas9 to introduce inversions into the genome of fission yeast. We either used two Cas9 plasmids, each with a unique single-guide RNA (sgRNA) sequence, carrying different selective markers, or used a single plasmid carrying both guides together. The plasmid or plasmids were transformed into fission yeast, together with repair fragments in which a sequence at the left side at the first breakpoint was combined with the reverse complement sequence from the left side of the second breakpoint, and similar for the right side (**Fig. 4a**). Transformants were then selected with one or two antibiotic markers, for the plasmid with two sgRNAs or the two plasmids with one sgRNA each, respectively. We introduced inversions from *his5* on the right side, to three different locations on the left side, generating inversions of 110kb, 225kb and 1.04Mb size. For each inversion, we used two 180bp long fragments containing 90bp from either side of the breakpoint as HR fragments. The obtained colonies were screened for *his*^-^ deficiency, followed by PCR for the inversion. The efficiency of obtaining the inversions differed between them. For example, while the shortest inversion was found in seven from eight screened colonies, the middle inversion was obtained in only two out of 24 colonies.

**Figure 4.**
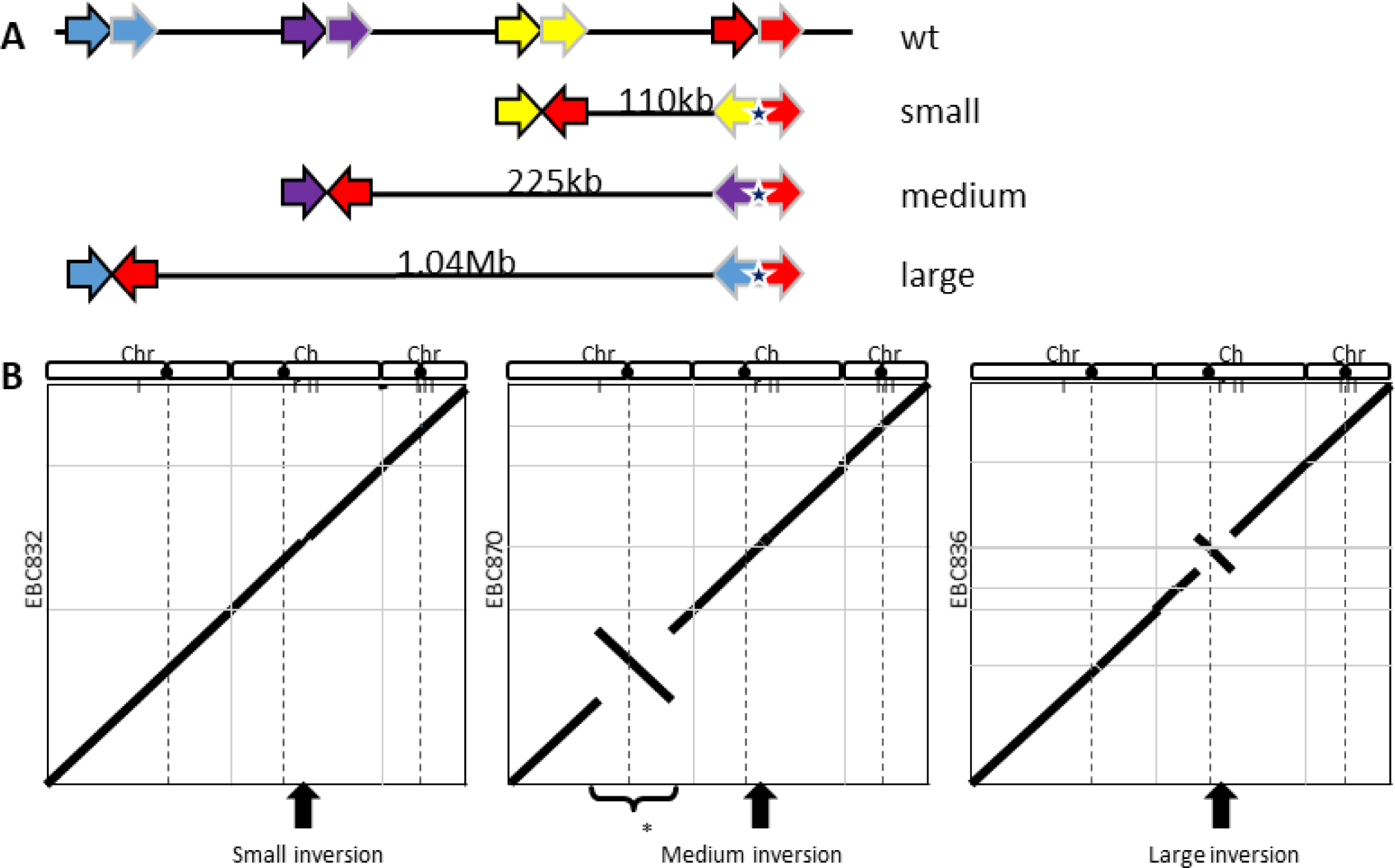
**A)** Overview of the three inversions (not to scale) and the HR fragments (colored arrows) used to guide repair of DSB to the inverted orientation. The right side of the left breakpoint is merged in reverse complementary direction to the right side of the right breakpoint (light outlined). The left side of the breakpoints are combined with the reverse complement of the left side of the right breakpoint (dark outline). The star indicates the location we introduced the natMX marker to track the inversion during crosses. **B)** Schematic dotplots showing the alignment of the three chromosomes of the reference on the x axis against the sorted contiguous contigs from the nanopore assemblies for three strains on the y-axis. The grey vertical lines demarcate the chromosomes as indicated at the top (black dot indicates centromere). The grey horizon lines indicate the ends of the contigs. The dashed lines indicate the location of the centromeres. The inversions generated in a single reaction using CRISPR/Cas9, are indicated with a large black arrow. The center plot shows next to the inversion described in the manuscript, the inversion indicated by the asterisk, which was generated by Hu et al. (2015) and crossed into this strain. See Figure SX for a dotplot containing all contigs per strain.

Using primers is fast, but for each inversion, four new 100bp long primers need to be designed and ordered. To overcome this limitation, we used the Golden Gate system to generate HR fragments using an (a-e) dummy to connect two target regions (**Fig. 1d**). Using the dummy elements, assembly of these HR fragments was efficient and greatly improved flexibility. Additionally, this gave us the possibility to introduce a selectable marker into the breakpoint, which allows tracking of the inversion. For this, we introduced a nourseothricin resistance marker into the right breakpoint. Furthermore, the flanking regions using Golden Gate are longer (∼500bp), which can increase HR repair efficiency. Verification of the inversions was performed by PCR, which showed again quite some variation in successful genomic manipulation. For example, two from sixteen colonies for the small inversion, two from twenty for the medium inversion and four from sixteen for the large inversion (**Fig. S3**).

We performed whole genome resequencing of some of the strains using nanopore long-read sequencing to analyse the breakpoints and asses off target effects. To test this method, we crossed a known inversion generated in (Hu et al. 2015) into one of our strains with the medium insertion. Visual inspection of the de-novo assembly of the genome showed the inversion in the location as expected without other large SV (**Fig. 4b**). To assess smaller SV, the reads were mapped to the reference genome and variant called with Sniffles2 (Smolka et al. 2022). These results showed no SV that were not present in our ancestral lab strain. Extraction of the breakpoints from the de-novo assemblies showed that the observed sequences at the inversion sites aligned perfectly with the designed regions, except for long homopolymer runs (**Supplementary data 1**), which is expected in nanopore data especially without polishing with short read data (Delahaye and Nicolas 2021).

## Discussion and outlook

The methods described here add to the available tools to use in fission yeast. Where much of the research so far has limited itself to genic modifications, our methods greatly improve manipulation of the genome at any location. Adding flexibility of elements that can be introduced at these locations, implementing the Golden Gate systems first introduced by Kakui et al. for fission yeast. Furthermore, by combining multiple sgRNA for CRISPR together with homologous sequences for DSB repair allows for precise production of large structural variants.

Easy modifications of the structure of the genome will be a great tool in studying a variety of topics. Fission yeast is an important model for the study of chromatin remodelling, because methylation of histone H3K9 and HP1 are well-conserved relative to animals, contrary to budding yeasts, which lacks these (Smirnova and McFarlane 2002; Oh et al. 2022). Recent research has shown the importance of the three-dimensional structure of the genome through topologically associated domains (TADs) on gene regulation, and chromatin boundaries and loops (Krijger and de Laat 2016). When structural variations can disrupt the normal boundaries and organization of TADs. For example, a deletion might remove sites of CTCC-binding factors and the cohesin complex, preventing it from forming the anchoring loop necessary for TAD formation (Zuin et al. 2014). Alternatively, duplications or translocations can introduce new binding sites that alter the interactions between TADs. Even though TADs appear to be limited in fission yeast (Mizuguchi et al. 2017; Gu et al. 2022), Hi-C sequencing experiments have shown strong interactions among centromeres and telomeres in *S. pombe*, as well as interactions between the mating-type region located on chromosome II and the right telomere from chromosome I (Mizuguchi et al. 2017). The same study showed interactions at smaller distances to occur along the genome however, their exact role is not clear (Benedetti et al. 2017).

Changes in the structure of the chromosomes further gives great opportunities to assess the function of inversions on reproductive isolation (Kirkpatrick 2010) and on the evolution of super-genes - clusters of tightly linked genes on a chromosome that are inherited together as a single genetic unit (Thompson and Jiggins 2014). Even though research in inversions has been performed for over a century, most experimental observations have been obtained using naturally occurring variants (Dubinin and Tiniakov 1946; Seoighe et al. 2000; Stevison et al. 2011; but see Grüneberg 1935). The consequences of inversions are that population level recombination in these regions is reduced and the regions diverge genetically from each other. The putative consequences of inversions could be confounded by genic differences occurring within the structural variants. Generating artificial structural variants will be able to separate these causes.

Many methods to perform genomic modifications are around and the yeast community is for a large part using a large number of well-established plasmids documented on pombenet.org maintained by the Forsburg lab. There is a trade-off with performing assemblies from PCR amplified fragments or cloning into a backbone. Cloning into a vector costs more and is more laborious than PCR. Nevertheless, we prefer cloning into a backbone, which is cheap and highly efficient using our *SmaI* backbone or the *BbsI* backbones for the target fragments. Additionally, due to the low mutation rate in *E. coli*, each element can be reused without the chance of PCR artefacts, avoiding the need to sequence each assembled plasmid. The golden gate system is a great and flexible system that could greatly benefit the fission yeast research community. The plasmids generated for this manuscript will be made available for any researchers interested on request and through the Japanese Yeast resource center. Since 2015, only few labs seem to have adopted the system introduced by Kakui et al., which after eight years has been cited only 21 times. We hope that with our extension, the toolbox becomes more useful for a larger group of researchers, who in turn can help increase the number of elements available.

## Methods

### Yeast strains, growth and media

All strains used in the experiments are derived from Leupold 689 strains (Jeffares 2018). The common strain used for transformations was EBC144, a strain derived from Klar & Miglio (Klar and Miglio 1986), that lacks the silent mating type regions *mat2,3*. Additionally, we used strain DY8531, which has an artificial inversion in Chromosome I (Hu et al. 2015), as a control for detecting inversions. An overview of the strains used are given in Supplementary Table S2. All strain propagation was performed on YES medium, crosses were performed on mating plates (EMM with NH_4_Cl replaced by 1g/L Glutamic acid), and we used PMG with appropriate supplements as minimal medium for all tests (Hayles and Nurse 2018). Tetrad dissections were performed on YES, using a MSM400 (Singer Instruments).

Yeast transformations were performed using the chemical synchronized cells protocol described in (Rodríguez-López et al. 2017) and selected on YES with appropriate antibiotics at 25µg/ml for nourseothricin and hygromycin and 50µg/ml for G418. We used ∼2µg CRISPR/Cas9 plasmid and for the HR fragment 10µl PCR product or 1µg linearized plasmid when *NotI* digested plasmid.

### Construction of plasmids

All plasmids are described in Supplementary table S3. In that table the primers used to generate these elements as well as the template DNA and references are given. The primers and their sequences are given in Supplementary Table S4. The receiving integration backbone plasmid pBN422 into which all elements are assembled, was generated by amplification with primers 743 & 744 – containing *Bsa*I sequences with f-h overhangs – of the ccdB and CmR chloramphenicol resistance genes from plasmid LIIc_F_1-2 (Binder et al. 2014) in which a *Sma*I restriction site was removed by iPCR. This fragment was integrated using *SmaI* integration into pUC57 as indicated in (Binder et al. 2014) and transformed in *E. coli* cells DB3.1 that are ccdA and selected for Chloramphenicol resistance. Donor backbone pBN111 was generated by introduction of a *SmaI* restriction site in a pCR-blunt-II-TOPO (Thermofisher) amplifying two fragments (primers 381 & 382 and 380 & 383) that were assembles using Gibson and transformed in *E. coli* strain DB3.1. This simultaneously removed a *SmaI* cut site present in the phleomycin resistance gene and introduced the required *SmaI* restriction site into the multicloning site. All plasmids and primers were designed using online tools in Benchling (benchling.com).

To improve efficiency of generating integration targets fragments, plasmids pBN195 and pBN196 were generated (**Fig. 1**). We amplified the ccdB/CmR fragment as for pBN422 using primer pairs 449&451 and 450&452, respectively, which were integrated into pBN111 using a *Sma*I integration (see below).

To generate episomal expression vectors, we PCR amplified *ars1* (Heyer et al. 1986) from pMZ379 with primers 439 & 440 and Gibson assembled (NEB) this with a pUC57 or pBN111 backbone amplified with 437 & 438. Into this backbone, a ccdB/CmR fragment was introduced using *Sma*I cut-ligation.

All *Sma*I restriction digestions were performed in a single reaction. A mastermix (1µl T4 ligase NEB, 0.3µl *Sma*I NEB, 1.5µl CutSmart buffer, 1.5µl T4 ligase buffer, 1µl pBN111/pBN103 [25ng/µl], 8.5µl H_2_O) was made to which 1µl of the diluted DNA fragment (0.7 ng/µl/kb) was added. This is incubated for 210 min at 25°C followed by 5 min at 50°C and 5 min at 80°C.

#### General Golden Gate assembly

For golden gate assembly with type II restriction enzymes *BbsI* (=*BpiI*) or *Bsa*I single tube reactions were performed. A mastermix (0.75µl T4 ligase NEB, 0.75µl *BsaI-HF V2* or *BbsI-HF* NEB, 1.5µl CutSmart buffer, 1.5µl T4 ligase buffer, 8.5µl H_2_O) was made to which 1µl of each element diluted to 13.3 ng/µl/kb was added, plus water to get to 15µl. This was incubated in a thermocycler using the following program: 5 min at 37°C, 30 cycles of 3min at 37°C and 3min at 16°C and finally 10 min at 37°C and 15 min at 80°C. Of this mix, 5µl was transformed into TOP10 *E. coli* and selected on LB with appropriate antibiotic. Assembled plasmids used in this study are given in Supplementary Table S5.

The fragment to generate a double CRISPR/Cas9 plasmids for the SpEDIT assembly method (Torres-Garcia et al. 2020) was generated by combining two fragments into backbone pBN196 using a *BbsI* Golden Gate assembly (as explained above). The first of these fragments contains the sgRNA sequence and the terminator region for the first guide, and the second the *tRNA* promoter and *HDV* ribozyme that will express the second guide RNA. The fragments were amplified by Phusion PCR using SpEDIT-plasmid pLSB-NAT as template with primers 741&742 (overhangs f-d) and 739&740 (overhangs d-a), respectively. Successful integration can be verified by screening colonies for lack of GFP signal.

To avoid potential non-homologous recombination with the inserted regions, we used sequences from other yeast species as shown by Li et al. (2019). We used sequences from *S. octosporus, S. japonicas or S. cryophilus* for a set of promoters (a-b) for which the *S. pombe* versions had been tested (Russell 1989; Iacovoni et al. 1999; Matsuyama et al. 2008; Verma et al. 2014; Wang et al. 2014). As template, DNA was used that was extracted from reference strains obtained from Yeast Stock Center Japan (strains FY21620, FY16936 and FY16937). Finally, we additional the fluorescent proteins *tagBFP*, *tdTomato*, *mCitrine* with linkers as (c-d) elements, and the three auxotrophic markers *ura4*, *arg3* and *his3* amplified from *S. octosporus* to be used as (g-e) elements.

### Integration vector construction (Levels 2, 3 and 4)

To ligate all the fragments, we diluted each to the same concentration of 20fmol/µl (i. e. ∼13.3 ng/kb/µl). Construction of Level 2 plasmids occurs by mixing together 1 µl of each element, with the backbone plasmid (pBN422, pBN213 or pBN231) in a single reaction with 0.75µl each of restriction enzyme *BsaI*-HF v2 and T4 ligase (both from New England Biolabs) and 1.5 µl each of their buffers (CutSmart Buffer and ligasion buffer including ATP). The mix was topped up to a total volume of 15µl with PCR grade water and cycled between 37°C (restriction) and 16°C (ligation) 30 times in a thermocycler to generate the new plasmid. After ligation 5µl of the mix was transformed in TOP10 *E. coli* competent cells using standard procedures (Sambrook et al. 1989) and were plated out on LB+amp. This selects against the donor plasmids, which have kanamycin resistance, and due to the ccdB sensitivity of the strains, any undigested receiver plasmid will induce death. The resulting colonies were multiplied in *E. coli* and isolated by mini-prep (Quick DNA Miniprep, Zymo Research). Colony PCR can be performed to assess proper construction using primers annealing to different donor elements. When large colonies are formed, these generally contain the correct constructs and this step can be omitted. Linearization of the integration fragment was performed either by Phusion PCR of the region using primers M13F and M13R, or by digestion of the regions using the *Not*I restriction sites that are present in the backbone directly flanking the f and h Golden Gate sites.

### Test of new elements

Promoters amplified from *S. octosporus* and *S. cryophylus* were assembled to express CFP together with an hphMX marker into a backbone containing the yeast autonomous replication sequence (see Table S4). The constructs were transformed into an *h^-S^* strain and selected using hygromycin (25ng/ml). The cells were grown in EMM to exponential phase and fluorescence signal was measured using flow cytometry on a BD Fortessa at CF FlowCyt at the Biomedical Center, Ludwig-Maximilians-Universität with identical settings and analysed in FlowJo for signal intensity. Similarly, complementation of strains was performed using prototrophic strains that were transformed using the *ura4* or *arg3* sequences clones from *S. octosporus*.

### Introduction and verification of inversions

We introduced three inversions of different sized into chromosome II of a *S. pombe* lab strain, using the same breakpoint on one side of the inversion, with different breakpoints on the other side. Each inversion was generated either with, or without introduction of the natMX resistance marker at the common breakpoint (**Fig. 4a**). Inversions without markers were generated either using 100bp long primers, which were extended to 180bp by PCR, or by golden gate assembly combining target donor elements that were directly connected using the dummy element (a-e). The latter generates fragments with longer homologous sequences. To add a selectable marker at one breakpoint of the inversion, we used golden gate, combining the targets with dummy elements (a-d and d-g) and the natMX marker sequence (g-e). Transformations were performed using synchronized cells, as described in Rodríguez-López et al. (2017) using two HR fragments, one for either side of the inversion, and two CRISPR/Cas9 plasmids each with one guide RNA sequence – one with kanMX and the other with hohMX as selectable markers – or using only hphMX when using a CRISPR/Cas9 plasmid with two guide RNAs.

Inversions were verified using yeast colony PCR of resistant transformants (Rodríguez-López et al. 2017), using Nippon FastGene Taq 2x Ready Mix. This kit is very efficient and can be used in a total volume of 10 µl per reaction, however, is sensitive to freeze-thaw cycles, so making aliquots is advised. Primers from either side of the inversion were used to test for no inversion, and one primer outside together with a primer inside of the inversion for positive controls. Only when a positive results for both sides of the inversion were observed did we consider the inversion to be correct.

To analyse the effect at the genome-level, whole-genome sequencing using Nanopore technology was performed. We isolated high-weight genomic DNA by growing 20ml cells to OD600 of 1.0 after which all cells were resuspended in 1M Sorbitol with 100 mM EDTA pH 8.0 and 100mg/ml Lallzyme MMX (Flor-Parra et al. 2014) and incubated for 30 min at 32°C with gentle agitation. The protoplast thus generated were spun down at 3500xg and the pellet resuspended in 1ml of Qiagen buffer G2 (800 mM guanidine HCl, 30 mM Tris·Cl, pH 8.0, 30 mM EDTA, pH 8.0, 5% Tween20, 0.5% Triton X-100) with 50 µl protinase K (>600 mAU/ml, Qiagen) and incubated at 37°C for 30min. This suspension was then extracted using a phenol-chloroform extraction method (Sambrook et al. 1989). After addition of ice cold isopropanol, the precipitated DNA was spooled on pipette tips, washed in 70% ethanol and resuspended in 100μl water. Library preps were performed for multiplexing and sequenced on a PromethION chip at Gene Center Munich.

The raw reads were basecalled with Dorado 0.3.0 duplex (Oxford Nanopore), demultiplexed and de novo assembled using Canu 2.0 with standard settings with genome size set to 14Mb. The inversion location was found by blasting the left or right side of the *his5* break point against these assemblies. The start and end coordinates with 5000bp on either side were then used to extract a fasta-file using *samtools faidx*. These were then aligned to the expected breakpoint sequences based on the reference genome (Wood et al. 2002). Whole genome comparisons were made with minimap2 (Li 2018) using D-Genies (Cabanettes and Klopp 2018). Additionally, we mapped the raw reads to the reference genome using minimap2 with the ‘ -x map-ont’ flag for nanopore data followed by transformation into bam, sorting and indexing using ‘samtools’ (Danecek et al. 2021), and analysed with Sniffles2 (Smolka et al. 2022).

## Acknowledgements

We thank the staff of the Core Facility Flow Cytometry, Biomedical Center, Ludwig-Maximilians-Universität, where flow cytometry was performed. We thank the undergraduate students in our lab over the last few years, especially Sebastian Schwartz and Chantal Krüger, who helped test and optimize the protocols, and perform cloning of the different elements.

---

**Figure S1.**
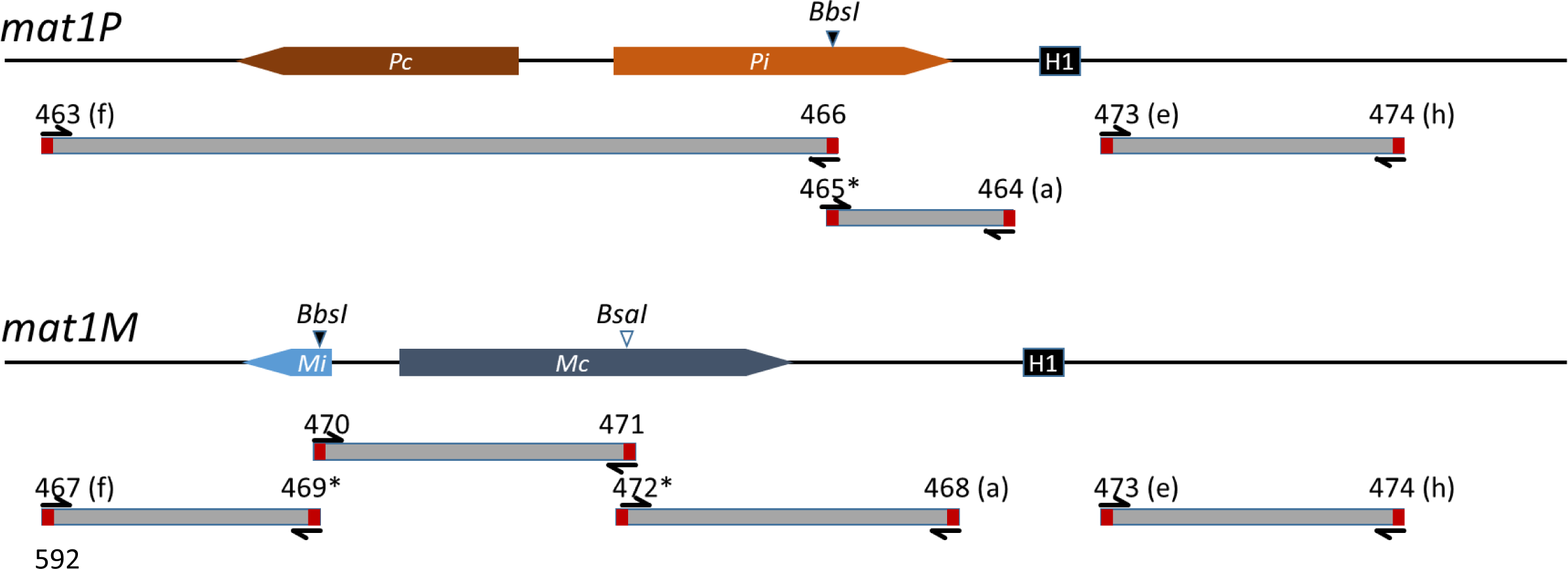
Overview of the construction of the target elements to introduce elements directly centromere proximate from mat1. The right side is the same for both *mat1P* and *mat1M*. The left side is constructed from two elements for *mat1P* and from three elements for *mat1M*. Each element is amplified from genomic DNA by proofreading PCR using the oligos indicated by the numbers. Each oligo has an overhang, indicated in red, which contains a *BbsI* restriction site that is removed during assembly, leaving overhangs complementing the element of the consecutive fragment, or with plasmids pBN195 (a-f) or pBN196 (e-h). The primers with * indicate a synonymous point mutation used to remove *BsaI* (filled triangles) or the *BbsI* (open triangle) in the wildtype sequence.

**Figure S2.**
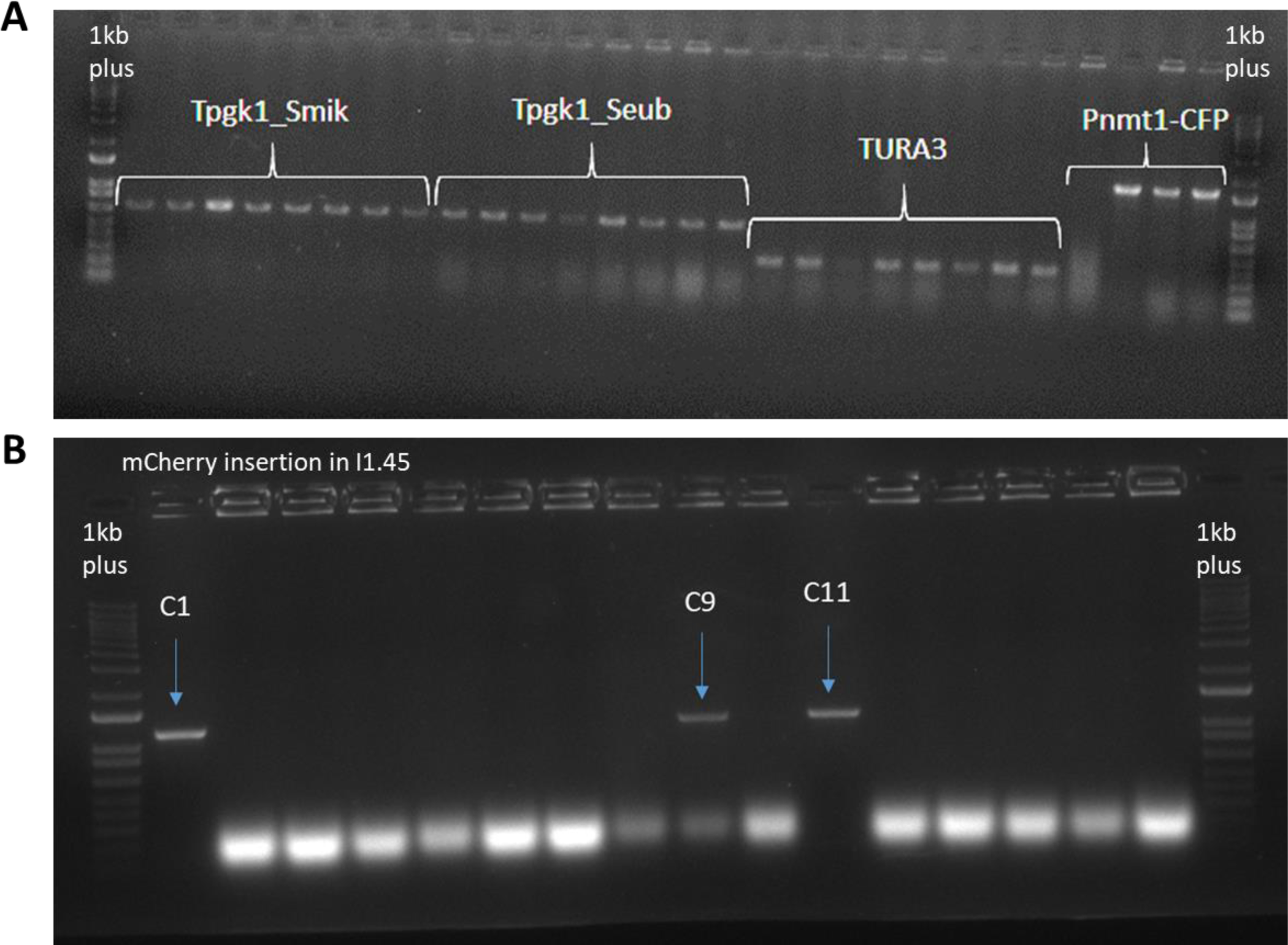
**A)** Example of cloning the terminator elements into pBN111 to generate level 1 plasmids. For each transformation all 8 colonies tested had the correct fragment. Cloning of a longer fragment containing the nmt1 promoter together with CFP worked in 3 from the 4 colonies tested. **B)** Example of yeast transformation results. For one transformation we tested 16 colonies and we had 3 positive results that showed the insertion in the intended place. The ladder used is Invitrogen 1kb-plus in both cases

**Figure S3.**
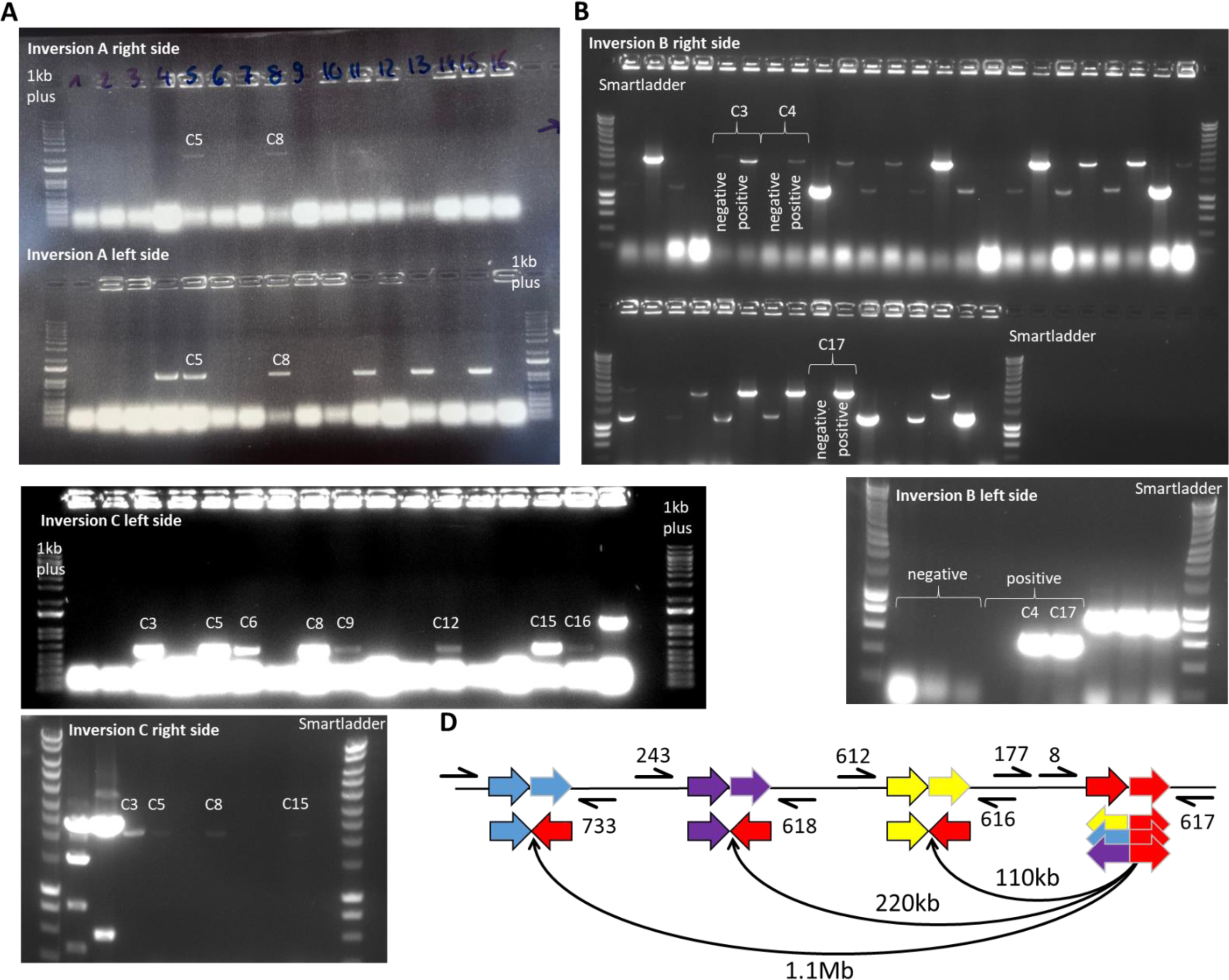
**A)** Results for confirmation of transformed inversion A (small). On top the positive results of the right side of the inversion (his5), checked with primers 616 and 617. On the bottom the positive colonies for the left side, tested with primers 612 and 177. From the tested colonies, we always chose the ones that had positive results for both sides, in this case C5 and C8. **B)** Results for inversion B (medium). In the top panel, every two lanes correspond to one tested colony. For each of them, the left lane tests for negative results (not inverted) with primers 7 and 8, and the right side tests for positive results (inverted), with primers 617 and 618. We chose the three colonies that do not give a band for a negative result and tested for the left side. On the bottom right panel, indicated the confirmation of inversion for colonies C4 and C17 with primers 243 and 8. **C)** Results for inversion C (large). On the top panel we tested presence of inversion on the left side with primers 732 and 8. With the positive results we tested on the right side with primers 733 and 617 and we show what colonies gave positive results (bottom panel). **D)** Overview of the three inversions (not to scale) and the HR fragments (colored arrows) used to guide repair of DSB to the inverted orientation. The setup and colours are the same as in figure 4a. Here we added an overview of the primers used for checking the presence of ineversions by PCR. Primers are represented in the DNA sequence before generating the inversions.

**Figure S4.**
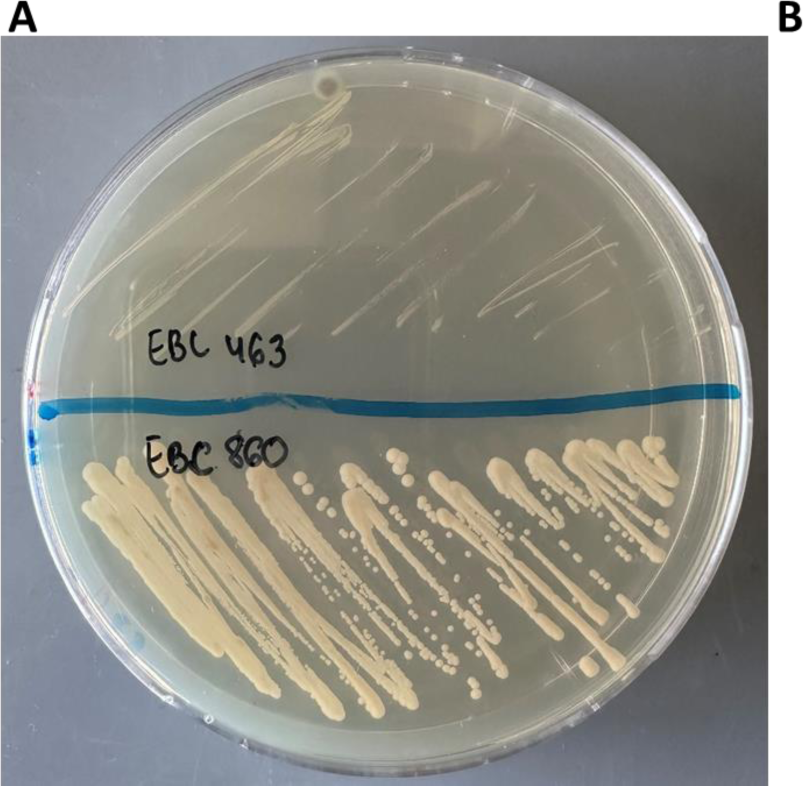
Overview of auxotrophic complementation of prototrophic *S. pombe* strains with markers *ura4, his3 and arg3* **A)** PMGuraDO plate with strain EBC463 (ura4 deficient) and strain EBC860, wich was transformed with pBN420. **B)** PMGhisDO Plate with strain EBC463 (his3-) on one side and EBC463 with his3 complemented by an expression vector (pCK005). **C)** PMGargDO plate with strain EBC463 (arg3-) on one side and EBC463 with arg3 complemented by an expression vector (pCK006). At the moment the transformations for the arg3 and his3 complementation are being performed, and proof of complementation will be updated in a future version

